# Disruption of thyroid endocrine in zebrafish exposed to BDE-209

**DOI:** 10.1101/176917

**Authors:** Dong Li, Xin Wang

**Affiliations:** Department of Environmental Toxicity, Third Hospital of Shanxi University

**Keywords:** BDE-209, HPT, Thyroid endocrine disruption, zebrafish

## Abstract

Polybrominated diphenyl ethers (PBDs) could adversely affect the thyroid endocrine system; previous studies report that BDE-209 has the potential effect on the fish thyroid endocrine system. In this study, we aimed to verify the bioconcentration and metabolism of BDE-209 in zebrafish. One day post-fertilization (dpf) zebrafish embryos were treated with different concentrations of BDE-209 (0, 0.01, 0.1 and 1 mg/L) until 10 dpf. BDE-209 was obviously accumulated in the zebrafish after 10 days exposure, and the metabolic products such as octa-BDE and nona-BDE were detected. After treated with BDE-209, the triiodthyronine (T3) and thyroxine (T4) levels were significantly decreased, suggesting that exposure to BDE-209 could disrupt the thyroid endocrine system in zebrafish. The transcriptional expression of genes involved in the hypothalamic-pituitary-thyroid (HPT) axis was altered. The mRNA expression levels of corticotrophin-releasing hormone (CRH) and thyroid-stimulating hormone (TSHβ) were significantly increased. The mRNA expression of pax8 and nkx2.1 which regulate thyroid development and synthesis were also increased. These data indicated that BDE-209 could disrupt the thyroid endocrine system in zebrafish, which could be assessed by hypothalamic-pituitary-thyroid axis.

## Background

Polybrominated diphenyl ethers (PBDEs) are some of the most popular brominated falme retardeants that are applied in many products worldwide [1-7]. BDE-209 has been reported having potential risk to the environment due to its persistence and may cause bioaccumulation in human and animals [8-14]. High concentrations of BDE-209 have been detected in sewage sludge and surface water, which might cause adversely effect on the aquatic organisms [10, 15-19]. Moreover, high levels of BDE-209 also have been detected in humans [3, 6, 8, 20, 21]. PBDEs might affect the thyroid endocrine systems due to their similar structures to the thyroid hormones [22-25]. Previous studies have been reported that BDE-209 can impair the thyroid endocrine activities in rodents [8, 26-28].

Hypothalamic-pituitary-thyroid (HPT) axis regulates the thyroid endocrine system in fish, and HPT axis controls the thyroid hormone dynamics through regulating their synthesis, secretion, transport and metabolism [29-33]. The potential effects of BDE-209 on thyroid endocrine disruption in fish have largely undefined.

In this study, we aimed to investigate the effects of BDE-209 exposure on the HPT axis in zebrafish and determine its effects on thyroid endocrine disruption. Zebrafish embryos were exposed to different concentrations of BDE-209 for 10 days, gene expression in the HPT axis were determined. Zebrafish larvae can efficiently absorbed and bioaccudmulated BDE-209, which led to developmental toxicity and thyroid endocrine impairment. Taken together, these results demonstrated that genes involved in HPT axis could be used for assessing the thyroid disruption effects of BDE-209 exposure in zebrafish.

## Materials and methods

### Chemicals

BDE-209 and DMSO were purchased from Sigma Aldrich. The BDE-209 stock solution (10 g/L) was prepared in DMSO as stored at ‒ 20 °C.

### Zebrafish husbandry and BDE-209 treatment

Wilde type zebrafish were maintained as previously described [34]. Embryos were treated with different concentration of BDE-209 solution (0, 0.01, 0.1 and 1 mg/L) from 1 to 10 dpf. Control group received 0.01% (v/v) DMSO. The BDE-209 solution was changed daily. Zebrafish were cultured at 28 ± 0.5 °C in a 14 h light/10 h dark cycle. After exposure, the zebrafish were collected for subsequent gene and protein analysis.

### Quantitative real-time polymerase chain reaction (qPCR)

Total RNA extraction and first strand cDNA synthesis were performed as described previously [35]. In brief, 50 zebrafish larvae per group were collected and homogenized for total RNA extraction by using RNeasy Mini Kit (Qiagen) and cDNA was synthesized using ProtoScript First Strand cDNA Synthesis Kit (NEB) follow manufacture’s instructions.

Quantitative PCR was performed using the SYBR Green Master Mix (Bio-Rad) and analyzed on a CFX96 Touch Real-Time PCR Detection System (Bio-Rad) [36]. The primer sequences were described somewhere else. The PCR conditions were: 95 °C, 5 min; 40 cycles of 95 °C for 30 s, 55 °C for 30 s, and 72 °C for 30 s.

### Thyroid hormone measurement

The extraction and measurement of thyroid hormone were carried out as previously described [37]. In brief, 100 zebrafish larvae of each group were collected and homogenized. After sonication, the samples were centrifuged at 6000 × g at 4 °C for 20 min, the supernatants were collected and used for thyroid hormone measurement.

### Statistical analysis

All data were shown as means ± SEM. The differences between the control and PDE-209 treated group were evaluated by one-way ANOVA followed by Tukey’s test by using SPSS 18.00. p< 0.05 was considered statistically significant.

## Results

### Toxicity of BDE-209 exposure to zebrafish development

Exposure to BDE-209 for 10 days has no obviously effect on the hatching and malformation rates in zebrafish. However, we observed the survival rates were significantly decreased in the BDE-209 treated group compared to the controls (Fig. 1).

**Figure 1.**
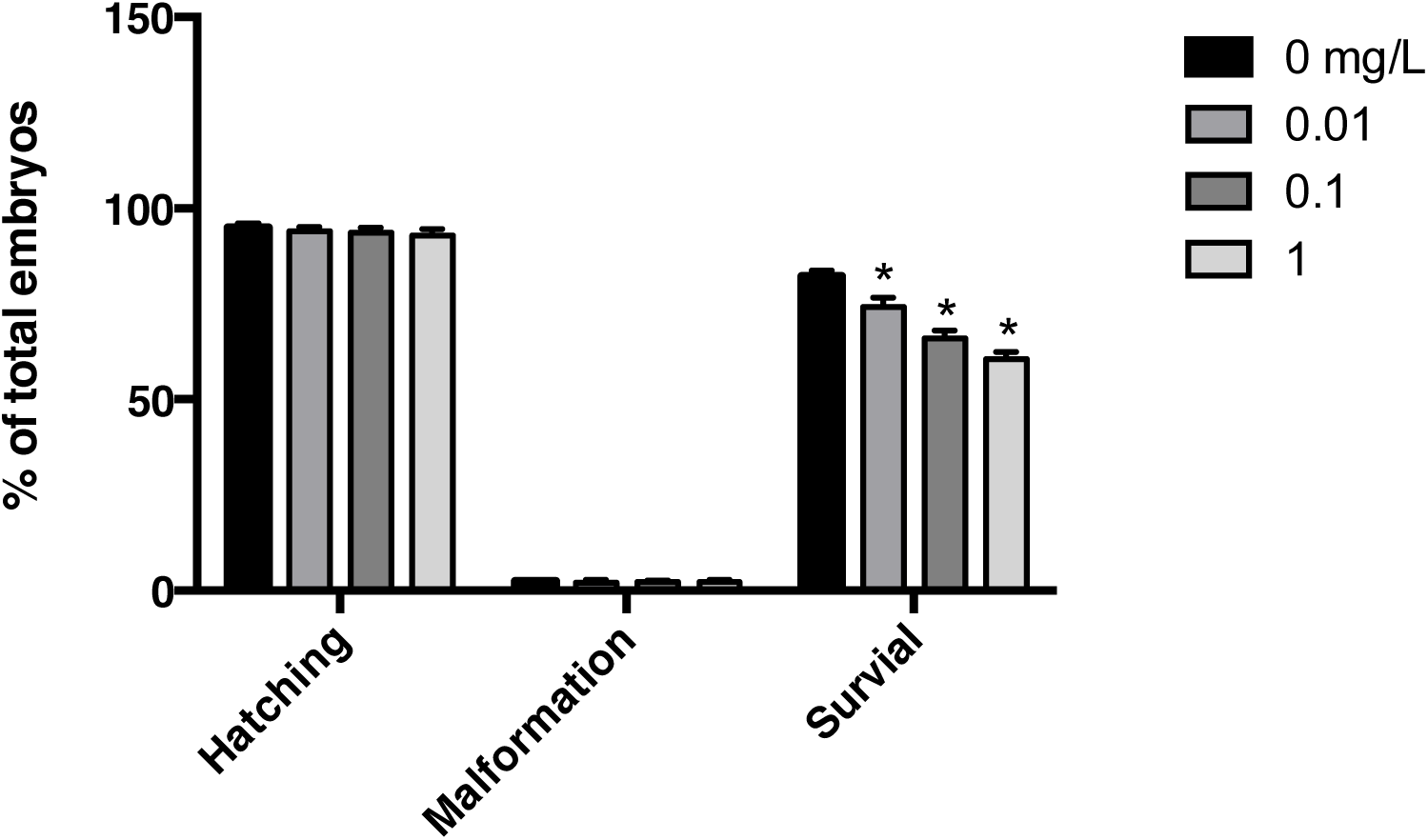
Developmental toxicity to zebrafish exposed to BDE-209.

### Gene transcription profile

We next examined the mRNA expression levels of genes regulate thyroid hormones transport, binding and metabolism. As shown in Fig. 2, the mRNA expression levels of CRH and TSHβ were significantly increased in a dose-dependent manner when treated with 0.01, 0.1 and 1 mg/L BDE-209. The transcriptional expression of genes involved in thyroid development was also significantly increased, such as pax8 and nkx2.1.

**Figure 2.**
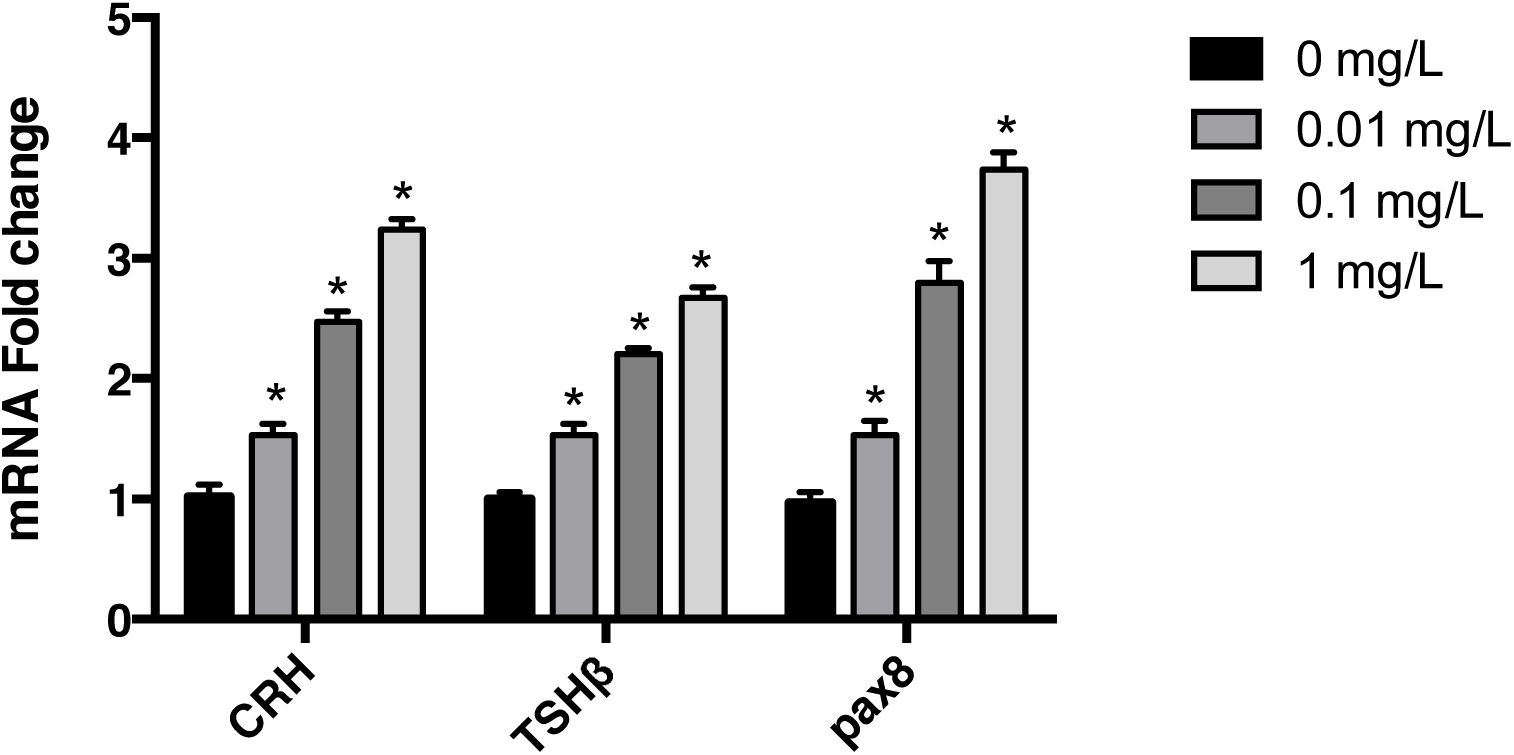
Gene expression in the HPT axis in zebrafish exposed to BDE-209.

### Thyroid hormones levels

The total thyroid hormones levels in zebrafish exposed to BDE-209 were examined after 10 days treatment. The T4 levels were reduced in the BDE-209 treated group (Fig. 3). However, the T3 levels in the treated groups were significant increased compared to the control group (Fig. 4).

**Figure 3.**
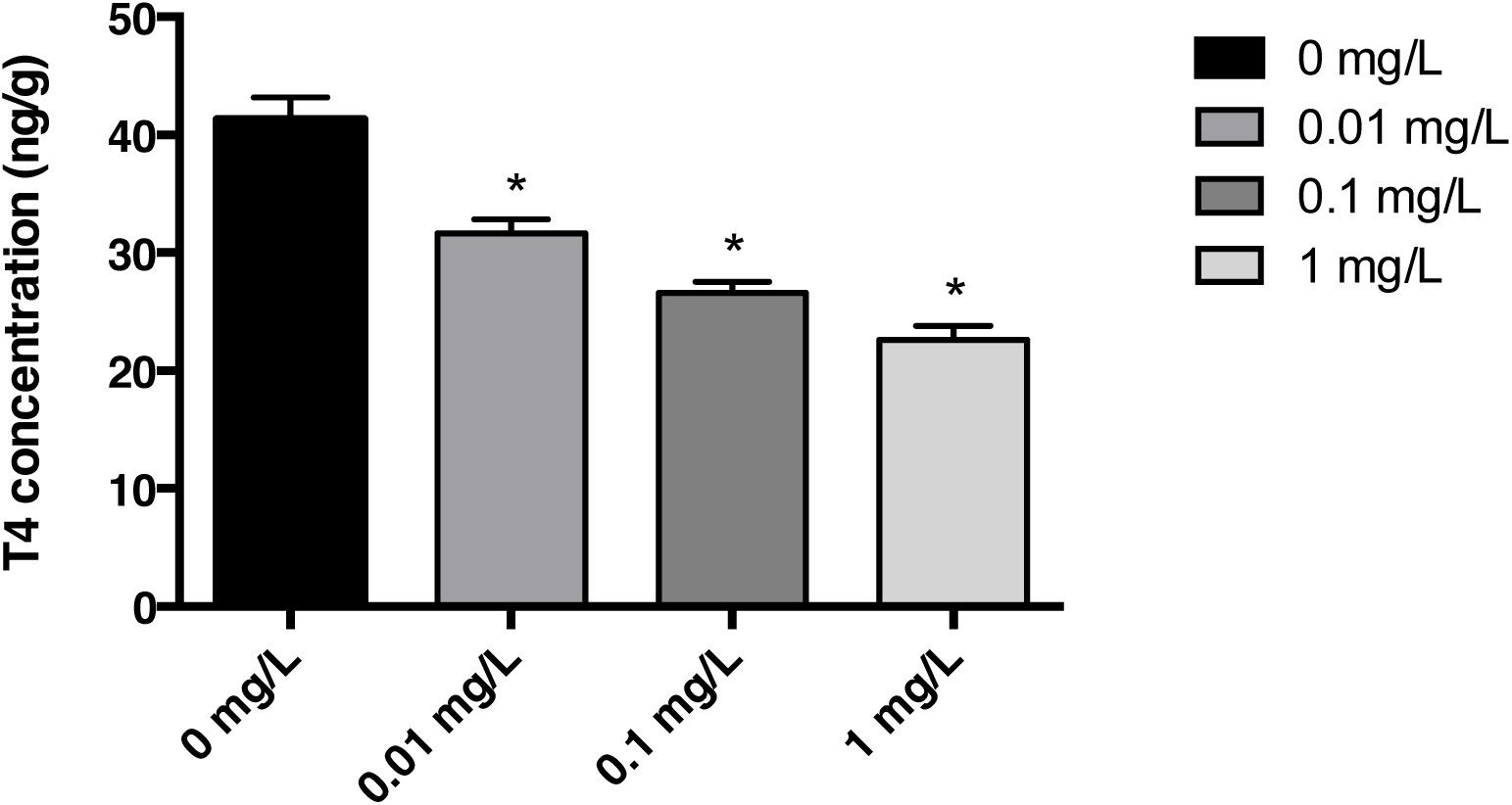
T4 levels in zebrafish exposed to BDE-209.

**Figure 4.**
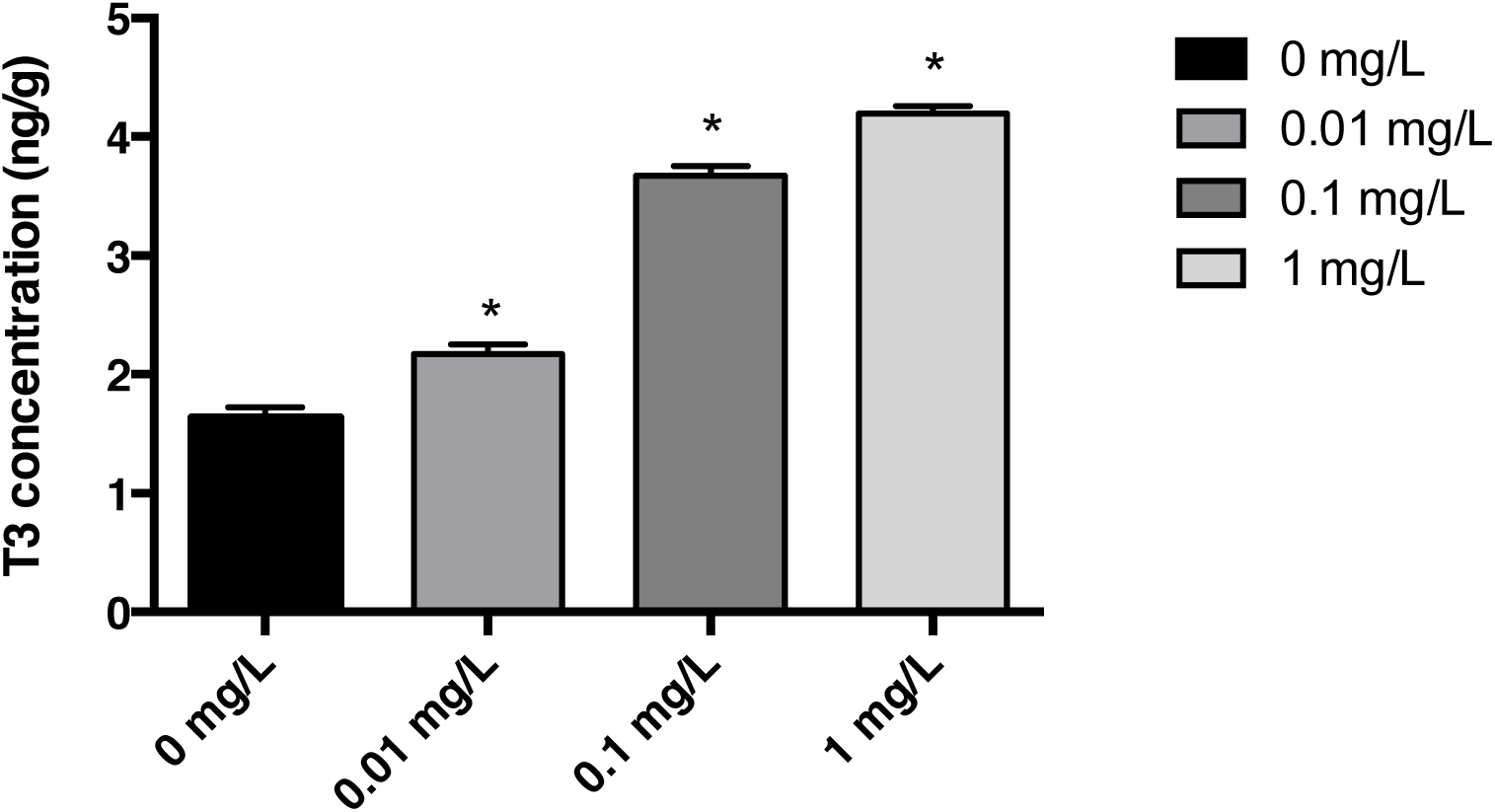
T3 levels in zebrafish exposed to BDE-209.

## Discussion

In this study, we evaluated the effect of BDE-209 on the thyroid endocrine disruption using zebrafish larvae. We found that BDE-209 could be bioconcentrated and metabolized in zebrafish larvae. And exposed to BDE-209 could affect the levels of T4 and T3 in zebrafish. These data suggested that we could use HPT axis to assess the effect of PBDE congeners on the thyroid endocrine impairment in zebrafish.

T4 levels were significantly reduced in BDE-209 treated embryos which confirmed the previous studies that BDE-209 impairs the thyroid in rodents and reduced the T4 levels [8, 38-40], however, T3 levels were increased in the BDE-209 treated embryos. These results suggested that PBDEs caused thyroid endocrine impairment by decreasing T4 contents, however, the levels of T3 might alter due to the treated time, species and doses of chemicals. These results indicated that the dose-dependent increases in the T3/T4 ratio was caused by the BDE-209 treatment that could interrupt the thyroid function.

The mRNA expression of CRH and TSHβ genes were significantly increased in the BDE-209 exposed groups, suggesting that the decreased T4 contents might form a negative feedback mechanism to regulate the relative gene expression [41]. The transcriptional expression of nkx2.1 and pax8 that were involved in thyroid development, were significantly increased in the PDE-209 exposed embryos.

In summary, we found that BDE-209 treatment can impair the thyroid hormones levels and the expression of genes involved in the HPT axis. Our results suggested that the HPT axis could be used for assessing the thyroid endocrine impairment of BDE-209.

